# Uncovering Hidden Data in the AMNH Marine Invertebrate Collections

**DOI:** 10.1101/2023.12.14.571719

**Authors:** A. Wilkinson, L. Berniker, C. Johnson, E. Rodriguez

## Abstract

Average global sea surface temperatures have been consistently higher during the past thirty years than any other time since reliable temperature recordings in 1880. Marine life is sufficiently adaptable to survive in a range of temperatures, but extended periods of temperature extremes can have a profound effect on the biology and behavior of aquatic organisms. Historic baseline data is often critical to understanding these changes in biology, distribution, migration, and population dynamics. Data from museum specimens can provide such baselines, but are often an overlooked resource, in part because museum collection data are not readily available. The American Museum of Natural History’s invertebrate zoology collection is home to an estimated 7.5 million marine specimens dating back to the mid-1800s. Although many of the specimens have been catalogued in a public online database, data from a large number of these specimens are not yet in digital form and are therefore only accessible to staff and researchers visiting or borrowing from the collection. The Museum’s Crustacea collection contains around 19,400 specimen lots which hold significant historic value, including lots from the New York Zoological Society’s Bermuda Oceanographic Expeditions.

The Bermuda Oceanographic Expeditions provide a rare opportunity to examine historic abundance and distribution of pelagic crustacea taxa that are currently experiencing extended periods of increased ocean temperatures. We synthesized data from the fluid-preserved specimens and from expedition field logs to compose a depth profile that compared the number of crustacean specimens, identified to family or lower taxonomic level, per depth. The results indicate that crustacean abundance and diversity was highest (4-7 families) between 500-1000 fathoms. Most of the specimens were diel-migratory shrimp from the family Sergestidae. We believe that this study provides a useful basis of historic sergestid distribution data that can be used to contextualize modern research in the warming Sargasso Sea.

## Introduction

The invertebrate zoology collections at the American Museum of Natural History (AMNH) contain an estimated 25 million specimens, 7.5 million of which are marine species. Curators and staff are currently working to digitize the majority of the non-mollusk marine invertebrate specimens supported by a National Science Foundation awarded Thematic Collections Network grant (digin-tcn.org/home). By photographing each specimen and recording all known identification and collection information, they hope to create a robust online database available to both researchers and the public.

Museum collections provide necessary context for modern research by establishing baselines to which new data can be compared (Boakes et al., 2010), and these baselines are particularly relevant in light of global warming. Establishing historical trends of crustacean distribution is particularly important because rising surface temperatures will likely affect the range of pelagic crustaceans. Piontkovski & Castellani (2009) found that zooplankton populations in the tropical region of the Atlantic decreased between 1950-2000, possibly due to the expansion of the “tropical belt” of warmer surface temperatures into higher latitudes and the unsuitably extreme increase in surface temperature in the tropics. It is possible that a similar phenomenon may be occurring in the Sargasso Sea; in fact, some species distribution models for North Atlantic zooplankton predict a general shift towards the poles as a result of global warming (Steinberg et al. 2012; Benedetti et al., 2021).

Global warming may also affect the vertical distribution of species within the water column, which has repercussions for the efficiency of the biological pump, or the process by which atmospheric carbon is gradually sequestered into the deep ocean (Steinberg & Landry, 2017; Devries et al., 2012). Recent studies have found that the biological pump is most efficient near the tropics (∼20°N-20°S) but has relatively high efficiency in the region around Bermuda (32°N) (DeVries et al., 2012).

In light of this, and with support from the NSF Research Experience for Undergraduates (REU) program, in the summer of 2023 we catalogued and digitized a subset of the AMNH crustacea collection. Our effort focused on crustaceans collected by the New York Zoological Society in the Sargasso Sea between 1929-1934 (geographic range of collection events presented in Figure 1). The New York Zoological Society was chartered in 1895 by a team of curators and researchers who hoped to inspire zoological research and encourage conservation efforts (New York Zoological Society, 1896). One of the society’s earliest members was prolific naturalist William Beebe, whose extensive contributions to marine science included pioneering in-situ research at depth in the Bathysphere, the first-ever manned submersible with a window. When trawling at extreme depths in the Hudson Valley, Beebe noticed that the collected specimens were extremely diverse, and wondered if such diversity was also evident in the region surrounding Bermuda (Beebe, 1931). This inspired him to lead the annual Bermuda Oceanographic Expeditions from which the specimens for this project were collected.

**Figure 1.**
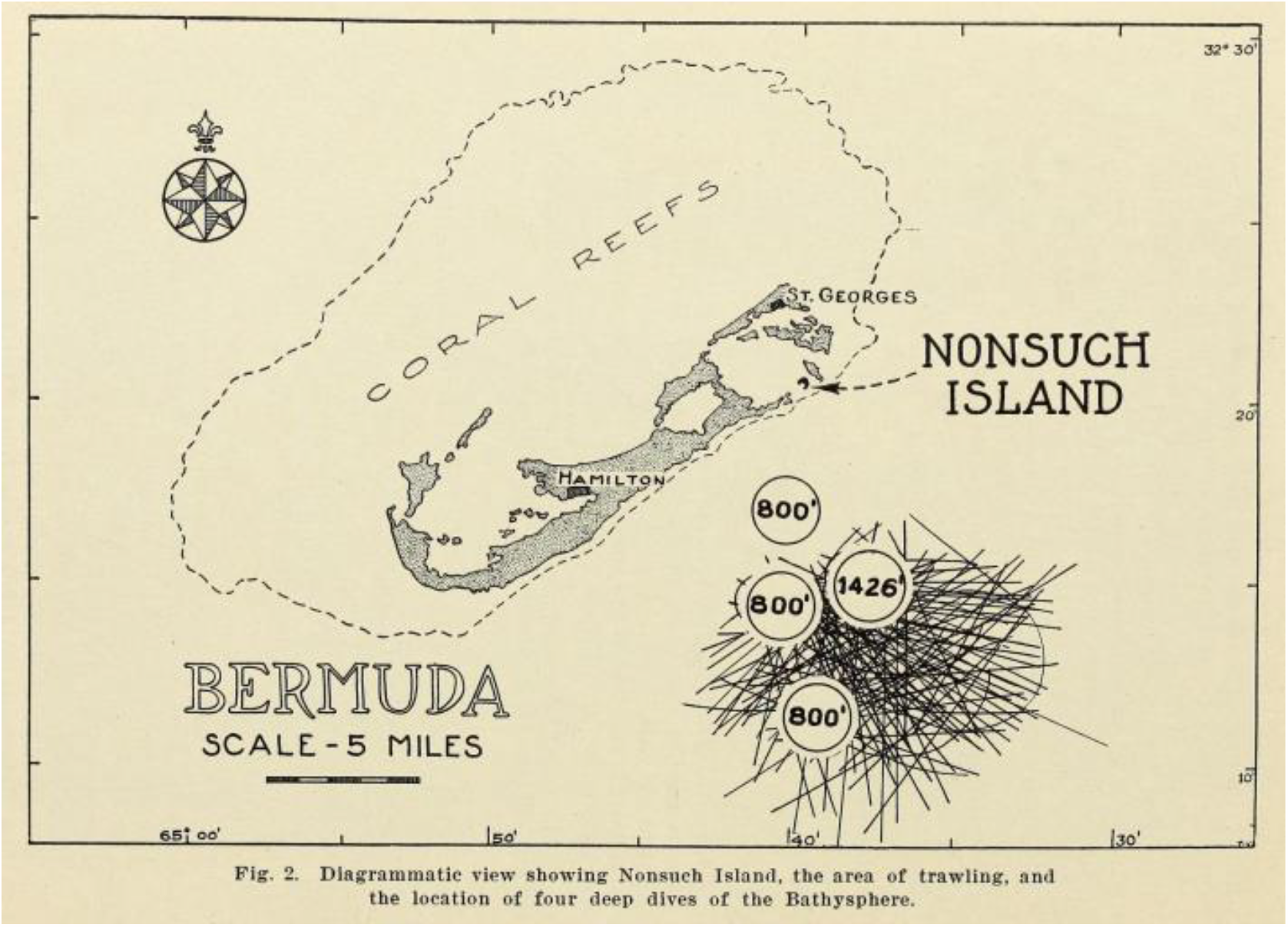
Diagrammatic view showing Nonsuch Island, the area of trawling, and the location of four deep dives of the Bathysphere. Adapted from Beebe (1931).

Unlike most collection events which serve as a single snapshot of a study region, the Bermuda Oceanographic Expeditions consisted of annual, six-month collection efforts in the same location over five years. Specimens were collected within one 8-mile radius site 9.5 miles SSE off the coast of Bermuda using nets at 100-fathom depth intervals ranging between 0-1200 fathoms. The in-depth perspective of the site over multiple years provided by this collection method makes the data from these expeditions uniquely useful for comparison to future surveys.

Specimens and records from the Bermuda Oceanographic Expeditions were acquired by the AMNH in batches between 1940 and 1972, and although many have been catalogued and databased the work is not yet complete. In addition to creating catalog records for the specimens, we endeavored to investigate trends in their abundance and distribution versus depth.

## Methods

### Rehousing

Most of the specimens were housed in their original glass jars, many of which were rubber-ringed bail jars. Specimens were removed from these jars and the original fluid (ethanol or formalin) was discarded. Specimens and any associated labels were then placed in new screw-top glass jars and filled with 70% ethanol.

**Figure 2.**
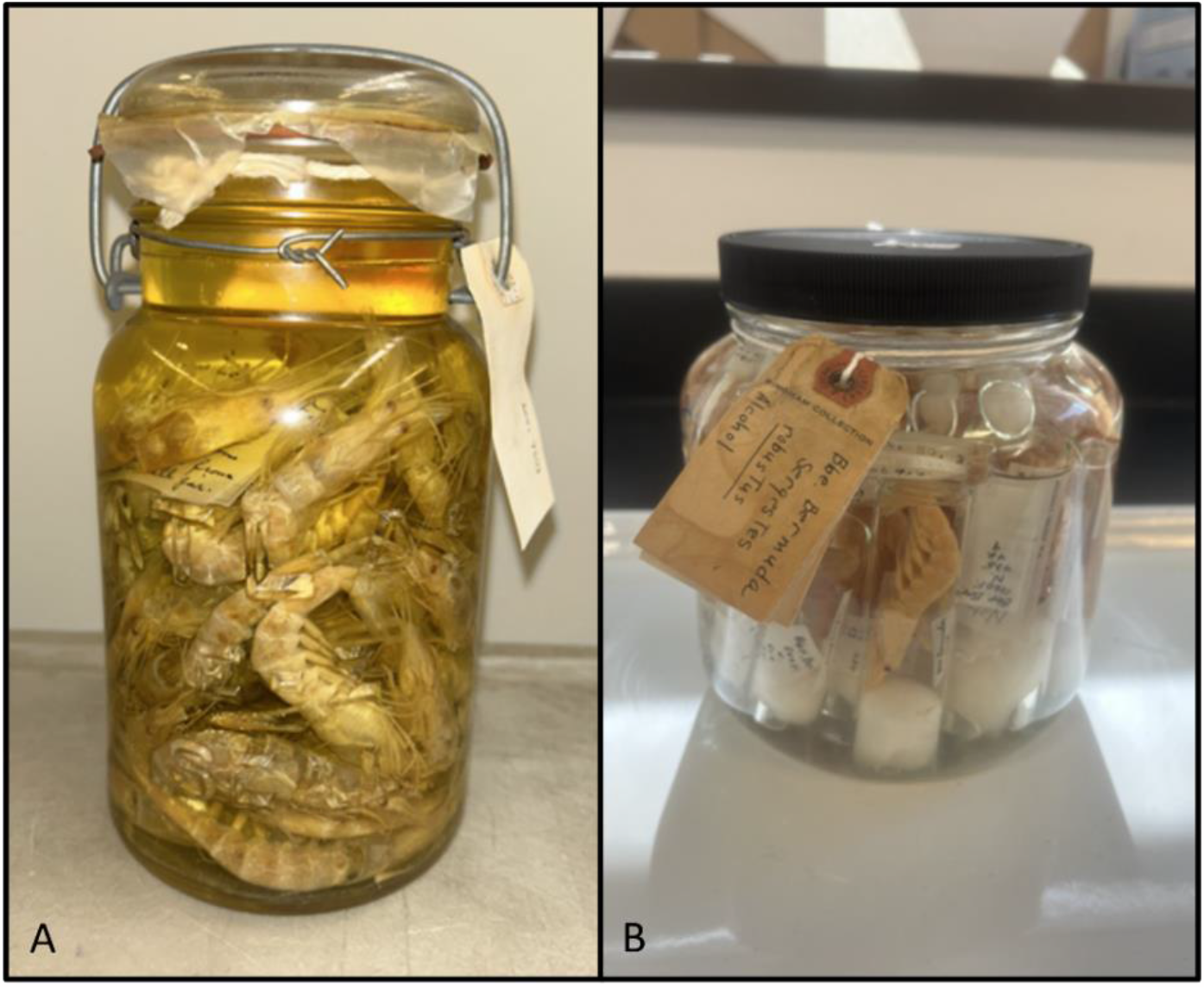
Fluid-preserved specimens before (A) and after (B) rehousing. A shows an example of specimens loose in a jar. B is an example of specimens in glass vials within a larger jar. Pictures by Anna Wilkinson.

### Photographing

Specimens were photographed using a CANON camera and lightbox. Images were taken via remote shooting through the program Digital Photo Professional. Specimen vials were placed on a clear plastic tray alongside a color calibration chart, ruler, “American Museum of Natural History” label, and any labels attached to the jar from which they came; specimens not contained in vials were placed directly on the tray. Each jar received a unique barcode, which was included in each specimen image from that jar. After every specimen was photographed, specimens were placed in their new jar, and the barcode was placed inside the jar. A sticky-back version of the barcode was attached to the top of the new jar to facilitate retrieving from the collection, and any labels from the original jar were tied around the new jar with twine.

### Digitizing

1. Uploading Images Images were renamed based on their included barcode number with BardecodeFiler (V2.7.2). New catalog records for each jar were created within the Museum’s database, Axiell EMu. Specimen images for each jar were attached to catalog records based on their corresponding barcode. Information written on labels from the original jars was manually transcribed into each catalog record.
2. Compiling Information Locality, collection data, and determination for each jar was gathered from publications (Beebe 1931, 1932, 1936) and from unpublished collection notes received with the material from the New York Zoological Society. All available information about each jar was transcribed into the corresponding record in EMu. For purposes of this study, only jars for which number of specimens, depth, and family were known were included in statistical analysis.

### Analyzing Data

Once all data was compiled, we analyzed family abundance and richness versus depth, as well as species abundance and richness within the family Sergestidae. All statistical tests were performed in R. A statistically significant difference between the number of nets at each depth was established via a Pearson’s Chi-squared test (p < 2.2e-16). This was used to revise Table 1 to account for sampling biases. For families found at multiple depths, an ANOVA test was performed to determine whether there was a significant difference in specimen abundance across different depths. It was found that there was no significant difference for any of the families (Aristaeomorpha: p=0.902; Benthesicymidae: p=0.545; Mysidae: p=1; Penaeidae: p=0.412; Sergestidae: p=0.981; Solenoceridae: p=0.488).

**Table 1.**
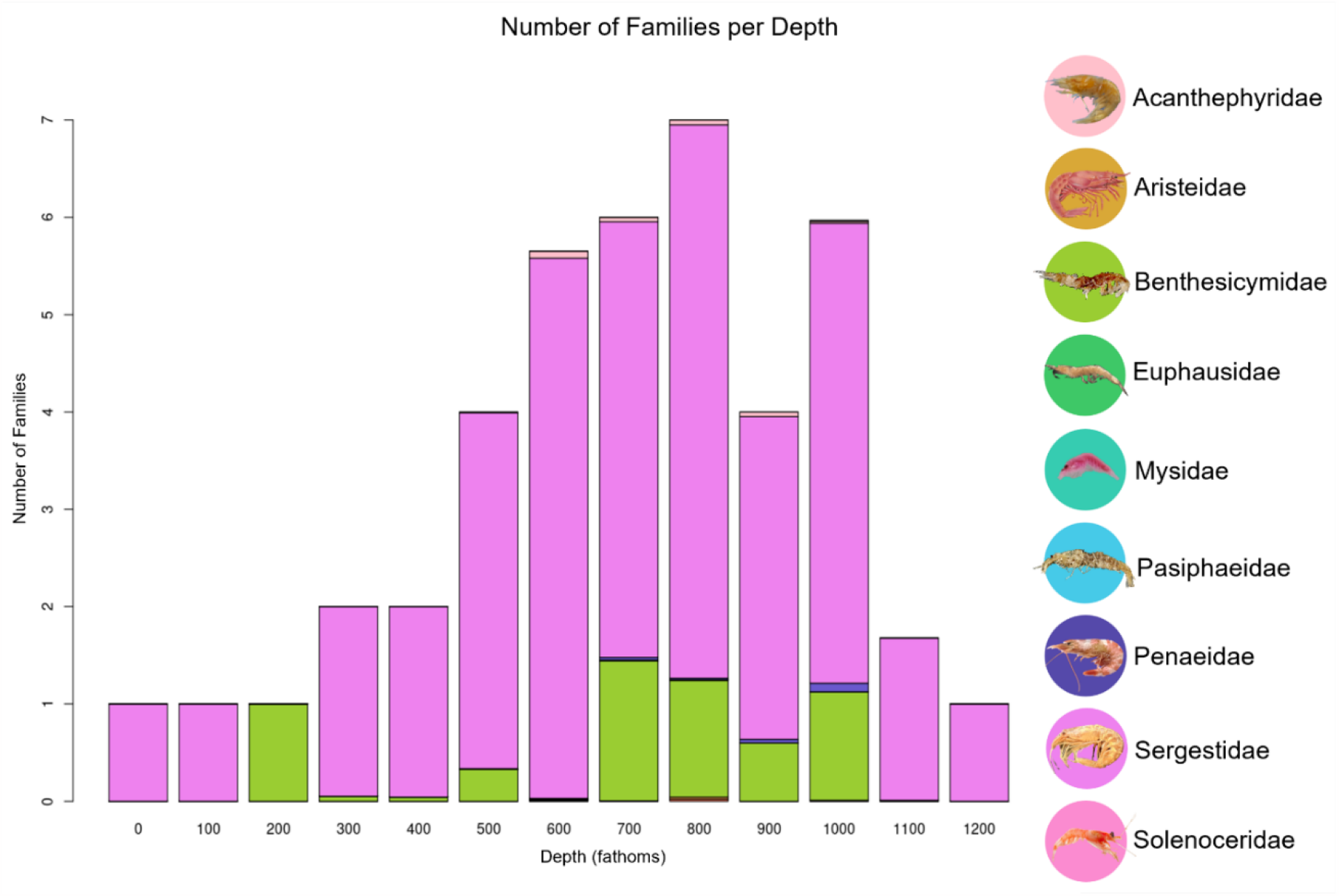
Family diversity is highest between 500-1000 fathoms, and Sergestidae is the most represented family. Colors correspond to the relative proportions of families per depth. Images of organisms are from the World Register of Marine Species (WoRMS).

All graphs were created in R.

## Results

In total, 114 new catalog records were created in Axiell EMu. These records contained 237 images and represented 9,498 specimens from 270 vials. The number of specimens is a low estimate, as vials for which the exact number of specimens was not known were recorded as a single specimen. Eight new locality records were created which correspond to depth ranges between 0-1200 fathoms at a point 9 ¼ miles SSE of Nonsuch Island, Bermuda. All these records are online and searchable through the Museum’s website (https://emu-prod.amnh.org/imulive/iz/iz.html).

The records created from this project represent 17% of the AMNH’s crustacean records from Bermuda and 21% of the crustacean records from the Bermuda Oceanographic Expeditions. 52% of the catalogued specimens were identified to species, 13% were identified to genus, 4% were identified to family, and 4% were identified to superfamily or higher. 27% of specimens in this collection were only identified as Crustacea and were therefore excluded from this analysis.

Our results indicate that crustacean richness and abundance in the Sargasso Sea near Bermuda between 1929-1934 were highest between 500-1000 fathoms (Tables 1 and 2). The largest proportion (82.2%) of these crustaceans was represented by the family Sergestidae (Decapoda: Dendrobranchiata) (Table 3).

**Table 2.**
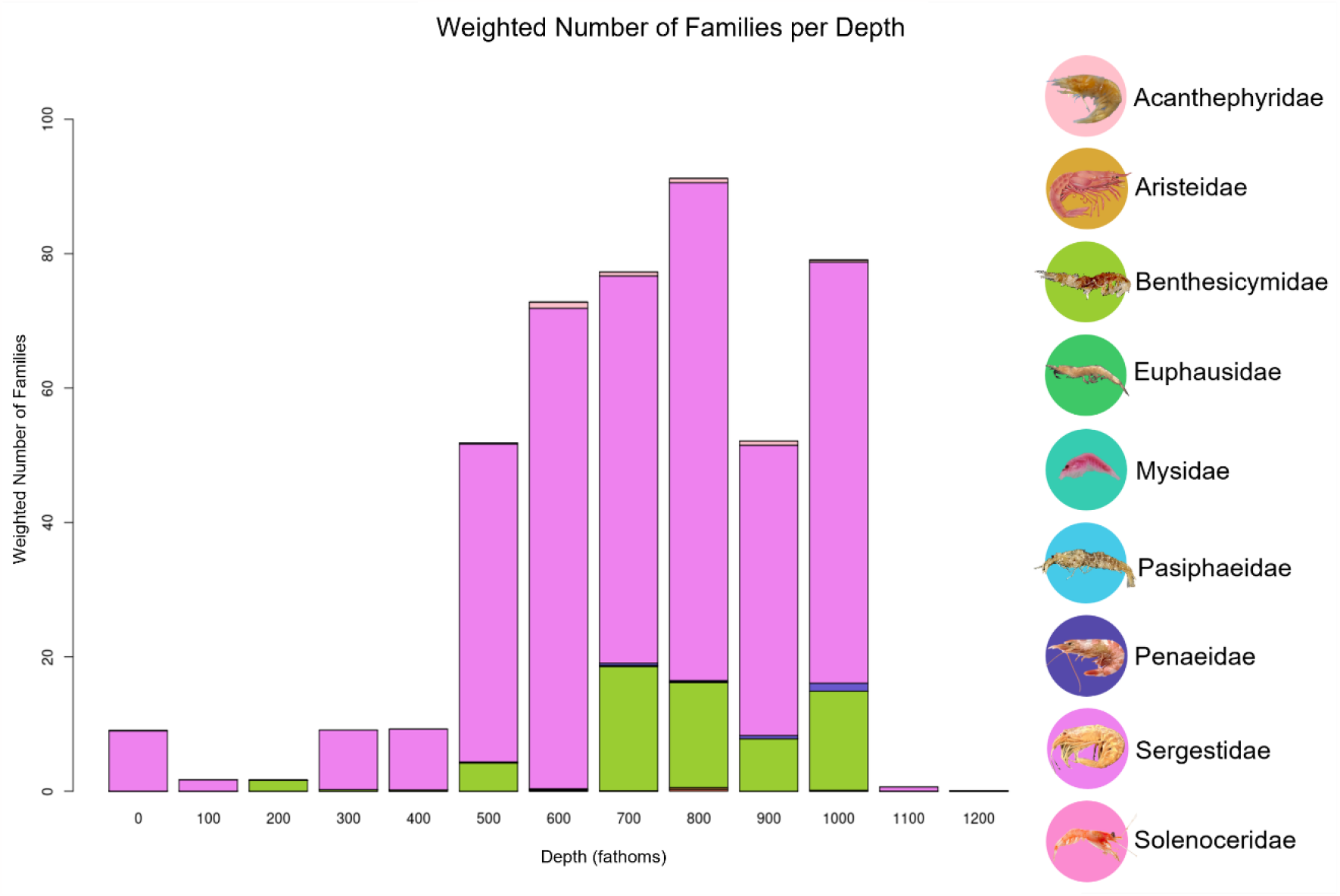
Weighted number of families versus depth shows a clear trend in high family diversity between 500-1000 fathoms. The relative percentages of nets at each depth were calculated, then values from Table 1 were multiplied by their respective net percentage value. Resultant values were multiplied by 100 for ease of visualization. Images of organisms are from the World Register of Marine Species (WoRMS).

**Table 3.**
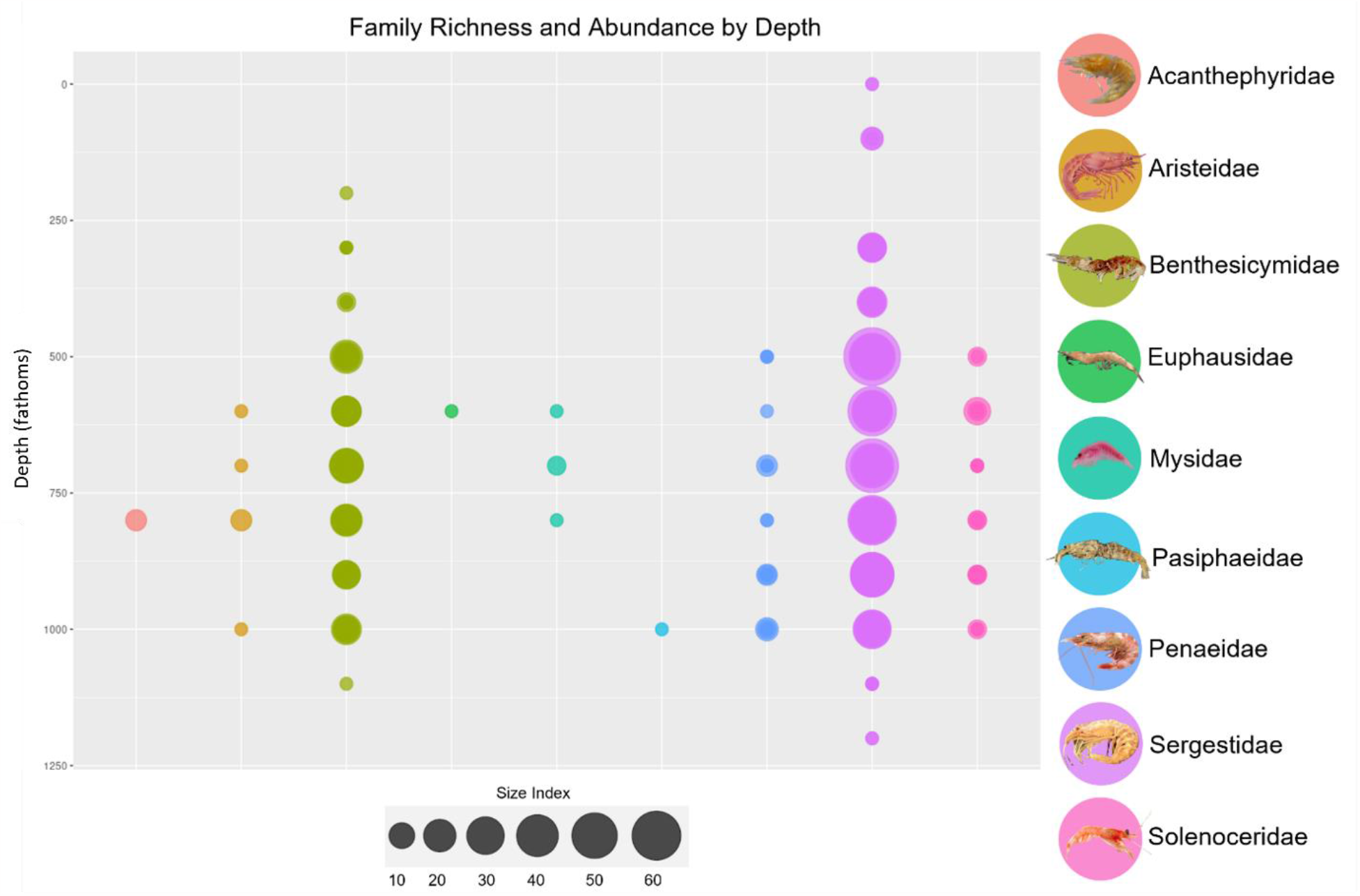
Family richness and abundance was highest between 500-1000 fathoms, and Sergestidae accounts for the largest number of specimens. Each point represents a locality at which specimens were found—point size indicates the number of specimens found at that point. Colors correspond to families. Depth is in fathoms. Images of organisms are from the World Register of Marine Species (WoRMS).

Within Sergestidae, the three most dominant species were all former members of the genus *Sergia* Stimpsom, 1860, which has now been split into eight genera (Vereshchaka et al., 2014). These species are: *Gardinerosergia splendens* Sund 1920, which represents 30.3% of the catalogued sergestid specimens, *Sergia remipes* Stimpson 1860, which represents 17.7%, and *Robustosergia robusta* Smith 1882, which represents 13.9%. The dominance of *Gardinerosergia splendens* (formerly *Sergia splendens*) amongst sergestids is consistent with modern research in the northern Atlantic Ocean (Vereshchaka, 1994; Ariza et al., 2015; Steinberg et al., 2020; Hine, 2022).

## Discussion

The depth profiles created from this data (Tables 3 and 4) provide meaningful context for recent studies on zooplankton distribution, particularly those associated with trends in global warming.

**Table 4.**
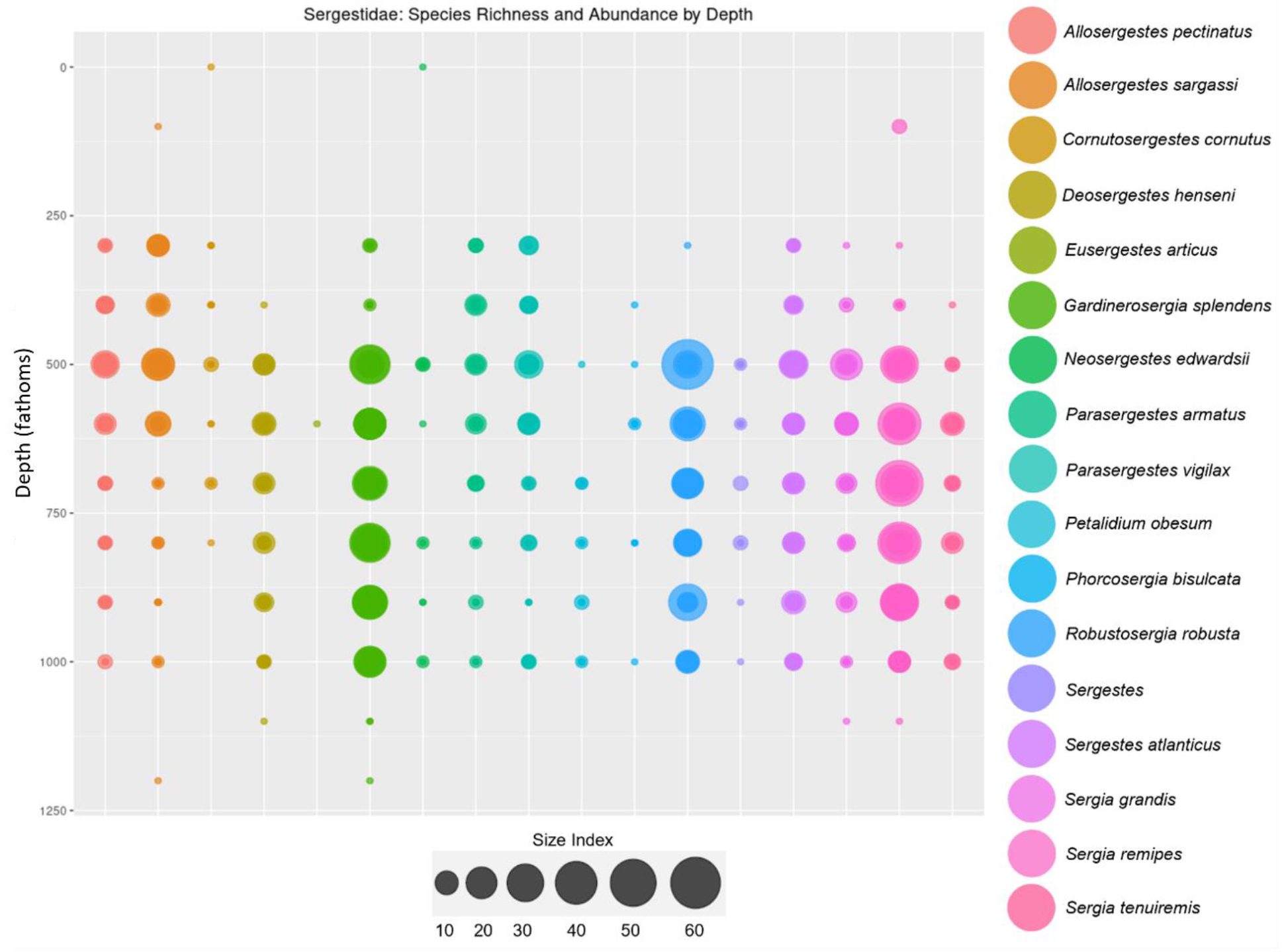
Species diversity within the Sergestidae family is high, and *Gardinerosergia splendens* is the most abundant Sergestid species. Colors correspond to species (or lowest identification). Depth is in fathoms. The column labeled “*Sergestes*” contains all specimens identified only to the genus *Sergestes*.

Steinberg et al. (2012) found that zooplankton biomass in the Sargasso Sea near Bermuda increased by 61% between 1994-2010 and was positively and significantly correlated with surface temperature. They posit that this increase was facilitated by an increase in phytoplankton biomass (an important food source for zooplankton) as climate-mediated increases in surface mixing and shrinking cold zones below the surface have supported primary production. They also noted that a large portion of the zooplankton were diel vertical migrators. Diel vertical migration describes a behavior in which individuals live at depth during the day to avoid visual predators, then migrate to the surface during the night to feed and spawn. This migration plays a key role in actively transporting nutrients between depths as part of the biological pump as well as providing a direct food source for meso- and bathy-pelagic carnivores (Flock & Hopkins, 1992). Diel migrators are therefore popular subjects for understanding the influence of global warming on bottom-up food chain control.

Across tropical and subtropical regions, sergestids are some of the most common diel migrators. The family *Sergestidae* consists of 97 species across six genera which are abundant in the Atlantic, Indian, and Central and West Pacific Oceans (Cardoso and Tavares, 2006; Vereshchaka et al., 2014). These decapod crustaceans range in size from approximately 10mm-50mm (Vereshchaka et al., 2014; Ariza et al., 2015).

Anthropogenic factors such as pollution may affect the vertical distribution of pelagic crustaceans. Most research on abundance and distribution of macrozooplankton has focused on shifts in latitudinal range rather than depth range, though Bos (2019) proposed that ingestion of microplastics may affect vertical migration patterns in crustaceans. Additionally, Hine (2022) established a detailed depth profile of sergestid species in the Gulf of Mexico to understand how distribution and biomass were affected by the Deepwater Horizon Oil Spill in 2010. Hine found that sergestid abundance significantly declined one year after the spill and continued to decline for six years. The depth profile from Hine’s study reveals that sergestid biomass was concentrated around the 0-600m range, which is roughly half as deep as the range of high biomass in the present Bermuda-based study.

Whether this difference is indicative of a larger-scale shift in depth distribution will depend on future research, as no detailed profile of crustacean depth distribution has been published for the Sargasso Sea since 1975 (Donaldson). It is our hope that the depth profiles presented in this paper will serve as a useful baseline to which future Bermuda profiles can be compared.

## Acknowledgements

We would like to thank the National Science Foundation for providing grant funding to digitize the AMNH marine invertebrate collection and for sponsoring the Research Experience for Undergraduates program through which this project was completed. We would also like to thank the American Museum of Natural History and the Richard Gilder Graduate School, particularly Dr. Jessica Ware and Asmeret Bekele, for supporting the REU program. Finally, we are grateful to Erin Willigan, without whose help and creativity this project could not have existed.

